# The Janelia Atalanta plasmids provide a simple and efficient CRISPR/Cas9-mediated homology directed repair platform for *Drosophila*

**DOI:** 10.1101/2023.06.17.545412

**Authors:** David L. Stern, Elizabeth Kim, Emily L. Behrman

## Abstract

Homology-directed repair (HDR) is a powerful tool for modifying genomes in precise ways to address many biological questions. Use of Clustered Regularly Interspersed Short Palindromic Repeats (CRISPR)-Cas9 induced targeted DNA double-strand breakage has substantially simplified use of homology-directed repair to introduce specific perturbations in *Drosophila*, but existing platforms for CRISPR-Cas9-mediated HDR in *Drosophila* involve multiple cloning steps and have low efficiency. To simplify cloning of HDR plasmids, we designed a new plasmid platform, the Janelia Atalanta (pJAT) series, that exploits recent advances in dsDNA synthesis to facilitate Gateway cloning of gRNA sequences and homology arms in one step. Surprisingly, the pJAT plasmids yielded considerably higher HDR efficiency (approximately 25%) than we have observed with other approaches. pJAT plasmids work in multiple *Drosophila* species and exhibited such high efficiency that previously impossible experiments in *Drosophila*, such as driving targeted chromosomal inversions, were made possible. We provide pJAT plasmids for a range of commonly performed experiments including targeted insertional mutagenesis, insertion of phiC31-mediated attP landing sites, generation of strains carrying a germ-line source of Cas9, and induction of chromosomal rearrangements. We also provide “empty” pJAT plasmids with multiple cloning sites to simplify construction of plasmids with new functionality. The pJAT platform is generic and may facilitate improved efficiency CRISPR-Cas9 HDR in a wide range of model and non-model organisms.

## Introduction

CRISPR-Cas9 induced mutagenesis has revolutionized experimental approaches in model and non-model organisms. DNA cuts induced by Cas9 or other enzymes^1, 2^ can be repaired by the non-homologous end-joining pathway to induce small deletions or cuts may be repaired by HDR if a homologous DNA template is present^3^. HDR provides the greatest power to manipulate gene function and to introduce new reagents into the germ line of experimental organisms, but CRISPR-Cas9 induced HDR works at low efficiency in many species.

HDR is normally implemented by introduction of three separate elements: Cas9 (or analogous enzyme), guide RNA, and homologous DNA. Usually, these three reagents are introduced as distinct components. In the most efficient systems, a source of gRNA and the homology arms are typically cloned into separate plasmids. Usually, plasmids containing homology arms are constructed by polymerase chain reaction (PCR) of homology arms from genomic DNA followed by cloning of arms together, often with a “payload” located between flanking homology arms. Generation of homology arms with PCR can be difficult for some target regions and cloning the homology arms with the payload also may have low efficiency, particularly when attempting to assemble multiple fragments.

We designed a new series of plasmids to simplify cloning for CRISPR-HDR experiments. The plasmid design includes two Gateway cloning sites for simultaneous introduction of long dsDNA fragments produced using new cost-effective DNA synthesis methods (Figure 1). The synthesized fragments incorporate homology arms between the *attL* Gateway cloning sites and one or both fragments includes a tRNA-sgRNA-tRNA array. We found that plasmids containing multiple copies of the standard Gateway negative selection marker, *ccdB*, were unstable^4^ and we therefore introduced a novel dual selection system to ensure efficient simultaneous cloning of two attL cassettes. We placed a U6 promoter sequence flanking one or both Gateway cloning sites to drive transcription of the tRNA-sgRNA-tRNA array, followed by a U6 terminator; this combination provides efficient precise release of sgRNA molecules^5^. Any payload can then be placed between the Gateway cloning sites, and we have generated a series of plasmids carrying some commonly used payloads.

**Figure 1.**
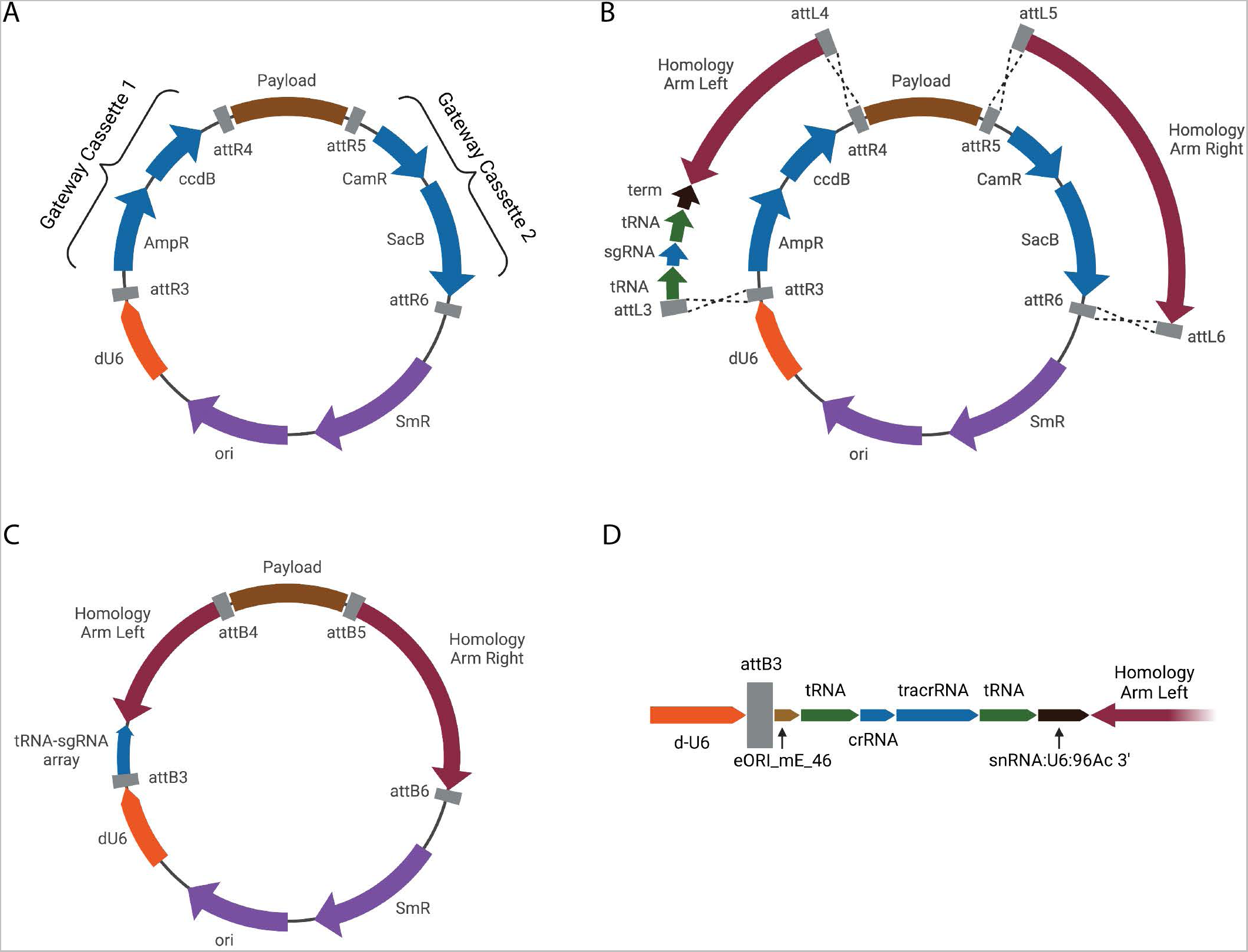
Design and use of the Janelia Atalanta plasmids. (A) Generic design of the Janelia Atalanta series. Each plasmid contains two Gateway compatible cloning sites, flanked on the left and right, respectively, by attR3-attR4 and attR5-attR6 sites. Each Gateway compatible cassette contains a different positive and negative selection marker. The negative selection markers *ccdB* and *SacB* can be selected simultaneously by cloning into *ccdB* sensitive cells and plating on minimal media containing a high concentration of sucrose. Each Atalanta plasmid contains one or two U6 promoters flanking the Gateway cassettes and a payload between the Gateway cassettes. (B) DNA fragments or plasmids containing paired attL3-attL4 and attL5-attL6 sites can be cloned into the based Atalanta plasmid simultaneously. In this example, synthetic DNA fragments containing left and right homology arms and a tRNA-sgRNA-tRNA array are cloned into the Gateway sites. The plasmid thus provides a source both of sgRNA and of homology arms for homology directed repair. (C) An example of an Atalanta plasmid after cloning of Gateway-compatible arms that is ready for injection into *Drosophila* embryos. Cas9 is provided either through co-injection of Cas9 mRNA or protein or by injection into embryos expressing Cas9 in the germline. (D) Detail of the components of a generic tRNA-sgRNA-tRNA region cloned downstream of the U6 promoter.

While we designed this plasmid for efficient cloning, we found that the plasmids yielded far higher HDR integration efficiencies on average than has been observed by us and others using other methods^3, 6, 7^. We have therefore named these plasmids the Janelia Atalanta (pJAT) series, after the mythological Greek heroine and exceptional archer (Supplementary Figure 1). The high efficiency of HDR with pJAT plasmids inspired us to attempt new kinds of experiments that have so far not been feasible in *Drosophila*, including introduction of chromosomal inversions.

The pJAT plasmids are modular and generic. One or two U6 promoter from any species can be introduced outside the Gateway cloning cassettes and any payload can be introduced between the cloning cassettes. pJAT plasmids may therefore be of value for HDR experiments in many model and non-model organisms.

## Results

### The pJAT series dual-Gateway cloning platform

Our initial aim was to build a plasmid platform that could utilize recent advances in DNA synthesis technology to simplify the cloning of reagents for CRISPR-Cas9 induced HDR. We found that some companies could synthesize long dsDNA fragments containing repetitive DNA attL sequences at the ends at a reasonable price (for example, 7 cents per bp from Twist Bioscience). This inspired us to design a dual-Gateway cloning platform for direct cloning of two fragments that each contain a homology arm (Gateway Cassettes 1 and 2; Figure 1A-C). To further reduce the number of plasmids that were required for experiments, we incorporated a tRNA-gRNA-tRNA design^5^ directly into the synthesized fragments (Figure 1B, D). Our initial attempts at building a dual-Gateway plasmid followed earlier examples and simply duplicated the standard *ccdB* negative selection gene in two separate attR-attR cassettes^8^. We found, however, that every plasmid recovered during cloning carried a mutation in one of the *ccdB* genes, suggesting that high *ccdB* dosage is deleterious to *E. coli*^4^. We therefore replaced one of the *ccdB* genes with a *sacB* gene, which allows negative selection by growing *E. coli* on plates containing high levels of sucrose^9^.

The pJAT plasmids contain one or two U6 promoters flanking the Gateway cloning cassettes or restriction sites that can be used to introduce new U6 promoters. We have generated pJAT plasmids containing multiple kinds of payloads, including (1) a germline source of Cas9, (2) attP landing sites, (3) marker genes, and (4) a split-GFP system to report on simultaneous HDR of two plasmids (Figure 2). All of the current pJAT plasmids utilize fluorescent reporter genes expressed in different anatomical regions^10^, which allow recombination of multiple reagents into a single wild-type fly without the use of balancer chromosomes. To facilitate introduction of novel payloads, we have generated one pJAT plasmid that contains a multiple cloning site for both standard ligation based cloning and Golden Gate cloning with BsaI or BbsI sites (Figure 2).

**Figure 2.**
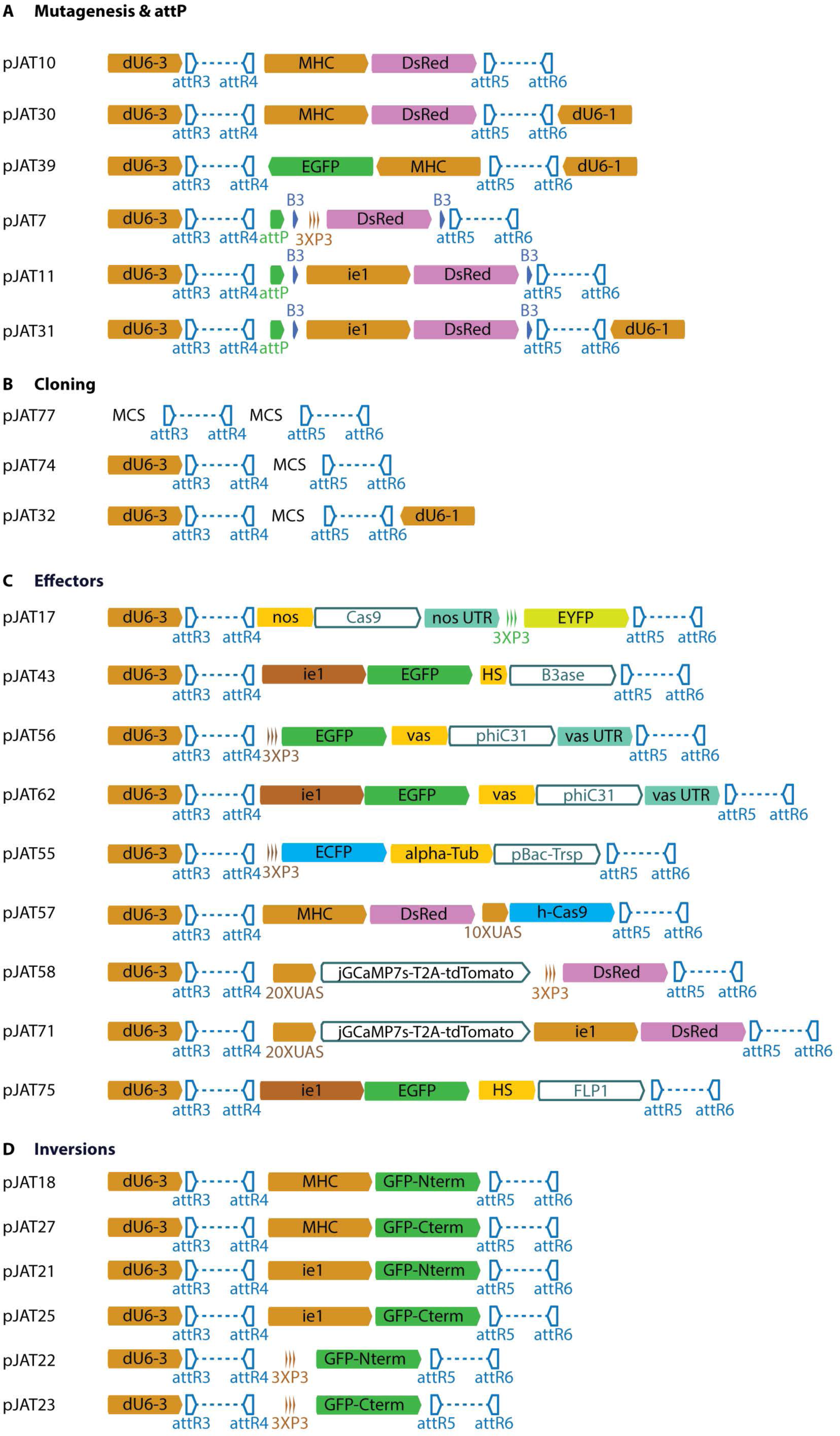
Payloads of Janelia Atalanta plasmids. (A) A set of plasmids containing fluorescent protein markers, attP sites, and removable fluorescent protein markers useful for targeted mutagenesis and generation of attP landing sites. (B) Plasmids to facilitate cloning of novel payloads. MCS=Multiple Cloning Site. (C) Plasmids carrying various effectors, including *nanos*-*Cas9* (pJAT17), B3 integrase under heat shock control (pJAT 43), and two plasmids expressing *vasa*-phiC31 integrase with different fluorescent protein markers (pJAT 56, pJAT 62). (D) Plasmids carrying split-*GFP* constructs under three alternative promoter systems for construction of chromosomal inversions.

For plasmid construction, homology arms were synthesized by Twist Bioscience with compatible attL sites and the company’s adaptor sequences are left on for three reasons: (1) synthesis sometimes fails without the adaptor sequences, probably because the repetitive attL sites are then located at the ends of the synthesized fragments, (2) keeping adaptors reduces synthesis cost; and (3) the adaptor sequences do not conflict with Gateway cloning. Synthesis of arms confers many benefits compared with PCR based construction of HDR plasmids, including the ability to include exogenous sequences, such as the tRNA-gRNA-tRNA array and other elements, as described further below, and the ability to introduce single base pair modifications. Synthesis by Twist Biosciences currently faces several limitations: (1) short homology arms are difficult to synthesize, probably because the repetitive attL sites are then positioned too close together; (2) the sequence cannot include more than 10 consecutive bp of the same nucleotide; (3) there are limitations on regions containing strong AT/GC bias and on repetitive sequences, which has prevented inclusion of multiple sgRNAs in a single synthesized fragment. Some of these synthesis challenges can be overcome by introducing innocuous point mutations, since HDR appears to be robust to considerable sequence variation relative to the target sequence. Cloning into pJAT plasmids requires standard Gateway cloning reagents and little hands-on time. A full protocol describing design of homology arms and all steps of cloning is provided at protocols.io.

### pJAT plasmids yield high efficiency HDR

Our initial experiments with pJAT plasmids yielded efficiencies of HDR of approximately 40-50%. We had previously only rarely observed such high efficiencies of HDR in *Drosophila* and so we systematically targeted many loci in *D. melanogaster* and found that we could successfully target all loci. We confirmed that all HDR products were integrated into the correct locations using TagMap^11^ with pJAT-specific primers (Materials and Methods). The average integration efficiency—that is, the fraction of fertile G0 injected eggs that yielded at least one positive event in the next generation—was approximately 25% for insertions into genes in which null mutations are homozygous viable and was more variable and lower on average for genes in which null mutations were homozygous lethal (Figure 3).

**Figure 3.**
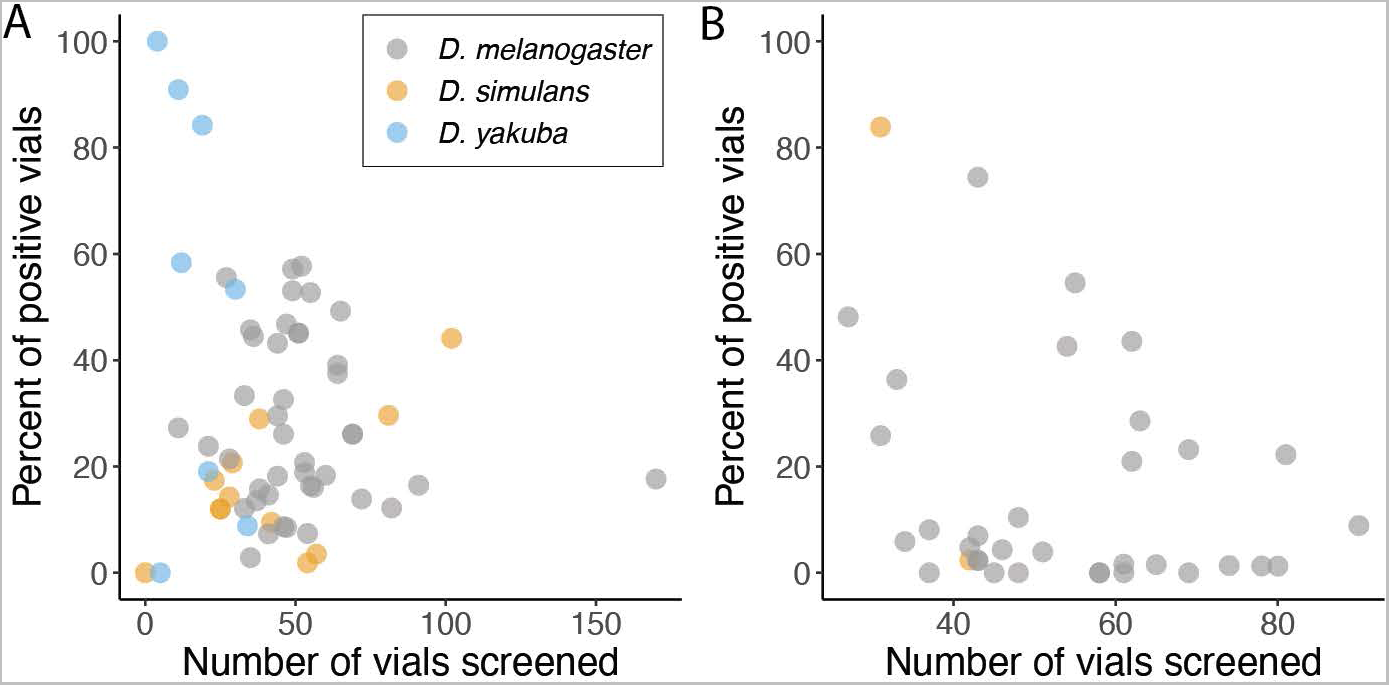
Integration efficiencies of Atalanta homology directed repair payloads into *Drosophila* species. The percent of positive events is plotted against the number of viable crosses screened. (A) Approximately 25% of injected animals producing fertile adults yielded integration events into genes categorized as “viable” on Flybase or into non-genic regions. No obvious difference was observed in integration efficiencies between the three *Drosophila* species *D. melanogaster*, *D. simulans*, and *D. yakuba*. (B,) Integration efficiency for payloads targeting genes categorized as “lethal” on Flybase (B).

Seven of the injections into lethal genes (shown as separate data points in Figure 3) yielded no events, but these were repeated injections using into only two loci. Despite these failures, other injections into these two loci yielded some events, though at low frequency. Thus, all genes were targetable with pJAT plasmids, but a few genes, and specifically a subset of those that are homozygous lethal, showed low efficiency. This low efficiency did not appear to result from the specific gRNA sequences used (data not shown) or the homology arm lengths (Supplementary Table 1). One possibility, which requires further investigation, is that the high efficiency of pJAT plasmids may induce homozygous null genotypes in many cells of the developing embryo, leading to lethality or sterility.

### Multiple features contribute to high efficiency of pJAT plasmids

To determine possible reasons for the high efficiency of HDR with pJAT plasmids, we systematically varied sections of the synthesized fragments. We identified two features that improved HDR efficiency. First, as has been documented previously^5^, flanking the sgRNA with tRNAs increased efficiency over co-injection with a sgRNA (Supplementary Figure 2). Surprisingly, a pJAT plasmid containing the sgRNA, but no flanking tRNAs, had similar efficiency to co-injecting with prepared sgRNA (Supplementary Figure 2), indicating that Cas9 can utilize a sgRNA even when it is embedded within flanking irrelevant RNA sequences, similar to results observed by others^7^. Second, use of intact dsDNA plasmids yielded higher efficiency HDR than plasmids that contained homology arms flanked by CRISPR-Cas9 cut sites (Supplementary Figure 3), which should linearize the plasmid DNA *in vivo* and which has been recommended previously^12^. It is possible that the relatively long homology arms used here improved efficiency, but we found that arms as short as 250 bp yielded similar efficiency as longer arms (Supplementary Table 1). We did not test shorter arms, since it is challenging to synthesize short arms with the long repeated attR sequences on both ends. The concentration of sgRNA produced from a single U6 promoter does not appear to be limiting, since synthesis of the same gRNA from two U6 promoters did not increase HDR efficiency (Supplementary Figure 4).

While pJAT plasmids provide sufficient gRNA for efficient HDR, we hypothesized that the quantity of Cas9 protein may be limiting when Cas9 is introduced exogenously, either as mRNA or protein. To test this idea, we built a pJAT plasmid that drives expression of a *Drosophila* codon optimized Cas9 mRNA from a germline specific *nanos* promoter with a *nanos* 3’UTR to localize Cas9 mRNA to the developing oocytes (Figure 2) and found that it increased HDR efficiency approximately 2-3 fold compared to co-injection with *Cas9* mRNA (Supplementary Table 2).

All of the experiments described above were performed by incubating injected embryos at 22°C after injection. Given the known temperature sensitivity of Cas9^13^, we performed a small test of incubation temperature by placing eggs at either 20°C, 22°C, or 25°C for 4 h after injection. We observed an increase in integration efficiency at higher temperatures (Supplementary Table 3), suggesting that the effect of temperature on CRISPR-Cas9 HDR in *Drosophila* should be investigated further.

Note that the efficiencies reported for many experiments shown in Figure 3 resulted from injections using Cas9 mRNA and with incubation at 22°C, which we now know are sub-optimal conditions. Thus, the average efficiencies reported here may underestimate the average efficiency using optimal conditions. The availability of a high-efficiency CRISPR-HDR platform that works in multiple species invites new kinds of experiments. We next illustrate two kinds of experiments that have been challenging to implement with previous technology.

### Scarless mutagenesis with pJAT plasmids

The use of synthesized homology arms and Gateway cloning can simplify the performance of scarless mutagenesis. One approach, which we have not explored, would be to insert a selectable marker at a target genome location and then, in a second set of injections, to replace the selectable marker with the intended genomic changes^14^. A second approach, which we illustrate here, is to place a selectable fluorescent marker between *piggyBac* transposon arms, which are included in the synthesized DNA fragments adjacent to homology arms that include the intended genomic changes^15^. In cases where the synthesized homology arms carry sites that differ from the genomic target, we have found that variable lengths of the synthesized homology arms, starting from the sites closest to the CRISPR cut site, are integrated into the genome. It is therefore best to place the intended genomic changes close to the *pBac* arms and to sequence the target region in HDR integrants to identify flies that carry the intended changes. The selectable marker, together with the *piggyBac* arms, is then removed by crossing flies to a strain expressing the *piggyBac* transposase gene (Figure 4). In the experiment performed here, we observed 58% integration efficiency (7 of 12 fertile injected G0 yielded integrants) into a site in *D. yakuba* and the *pBac* sequence was efficiently removed from multiple lines. This approach allowed efficient recovery of two scarless individual nucleotide changes at the targeted locus (Supplementary Figure 5).

**Figure 4.**
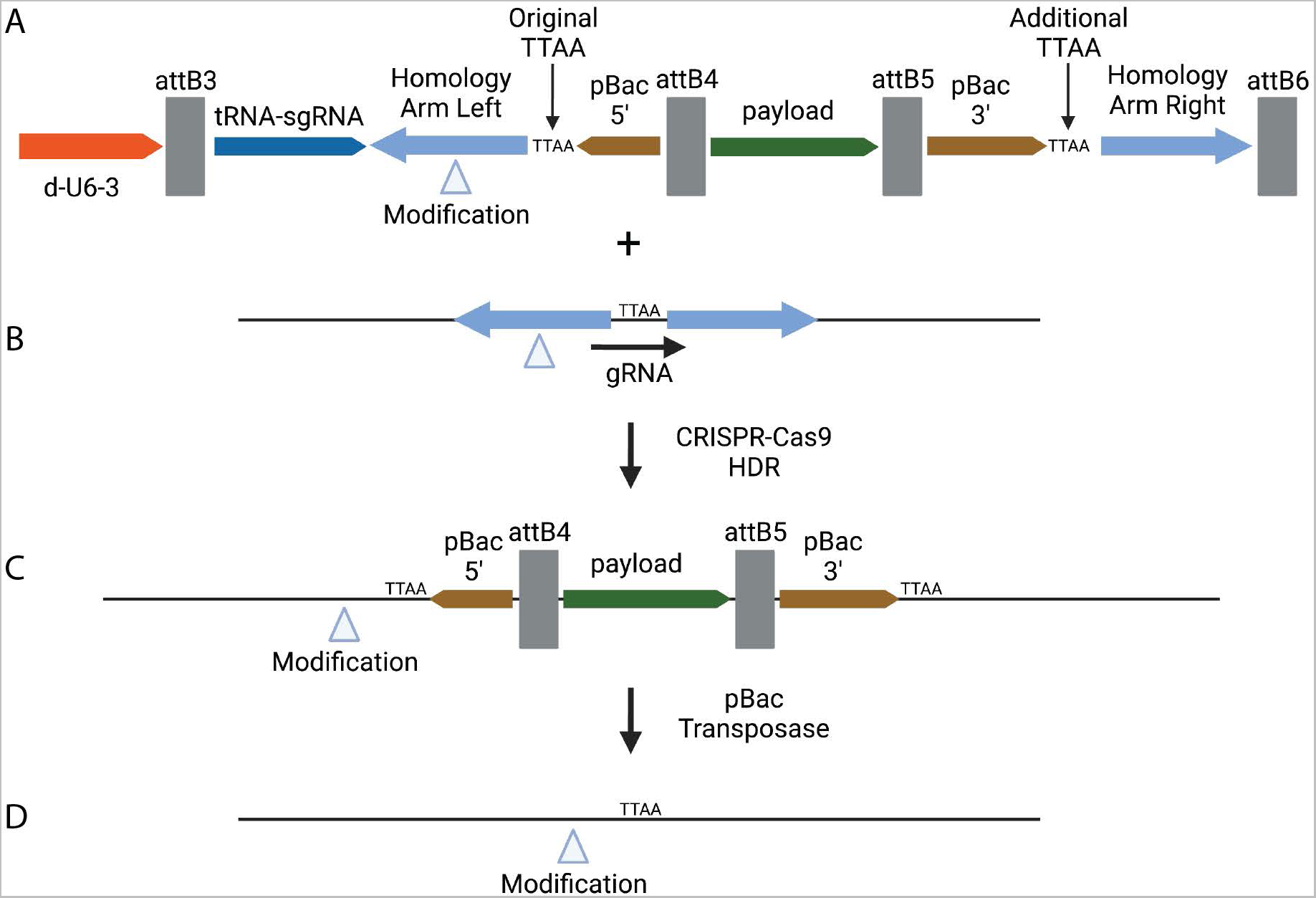
Schematic representation of scarless genome modification using pJAT plasmid. (A, B) Diagram of a pJAT plasmid illustrating the location of the synthesized *pBac* transposon 5’ and 3’ arms internal to the synthesized homology arms (A). The gRNA, included within the tRNA-gRNA-tRNA array on the left homology arm, targets a genomic site that includes a TTAA motif (B), the native target site for the *pBac* transposon. The synthesized right homology arm includes an additional TTAA sequence between the 3’ *pBac* sequence and the right homology arm. Intended modifications to the genomic sequence can be included in either or both homology arms. (C) Representation of the genomic sequence after integration of the pJAT plasmid. Flies containing the intended genomic modification(s) can be identified by PCR and sequencing. (D) After exposure to a source of *pBac* transposase, flies can be recovered that have lost the reporter gene and contain the intended scarless genome modification.

### pJAT plasmids can direct one-step targeted chromosomal inversions

The HDR efficiency of pJAT plasmids was considerably higher than we have observed using other approaches and this high efficiency emboldened us to try more complex experiments. One experiment that would be of considerable value, especially for studies of non-*melanogaster Drosophila* species, would be the ability to generate chromosomal inversions that could subsequently be used to balance deleterious alleles^16^. This experiment requires efficient cutting at two chromosomal sites followed by inversion of the intervening sequence (Figure 5A). To encourage production of the inversion event, we designed a pair of pJAT plasmids where each plasmid carried homology arms oriented in the same direction as the intended inverted chromosomal region (Figure 5B). To simplify detection of simultaneous integration of both plasmids, potentially indicative of an inversion, we built three pairs of pJAT plasmids expressing split-GFP in either the eyes, the thorax, or the abdomen (Figure 2).

**Figure 5.**
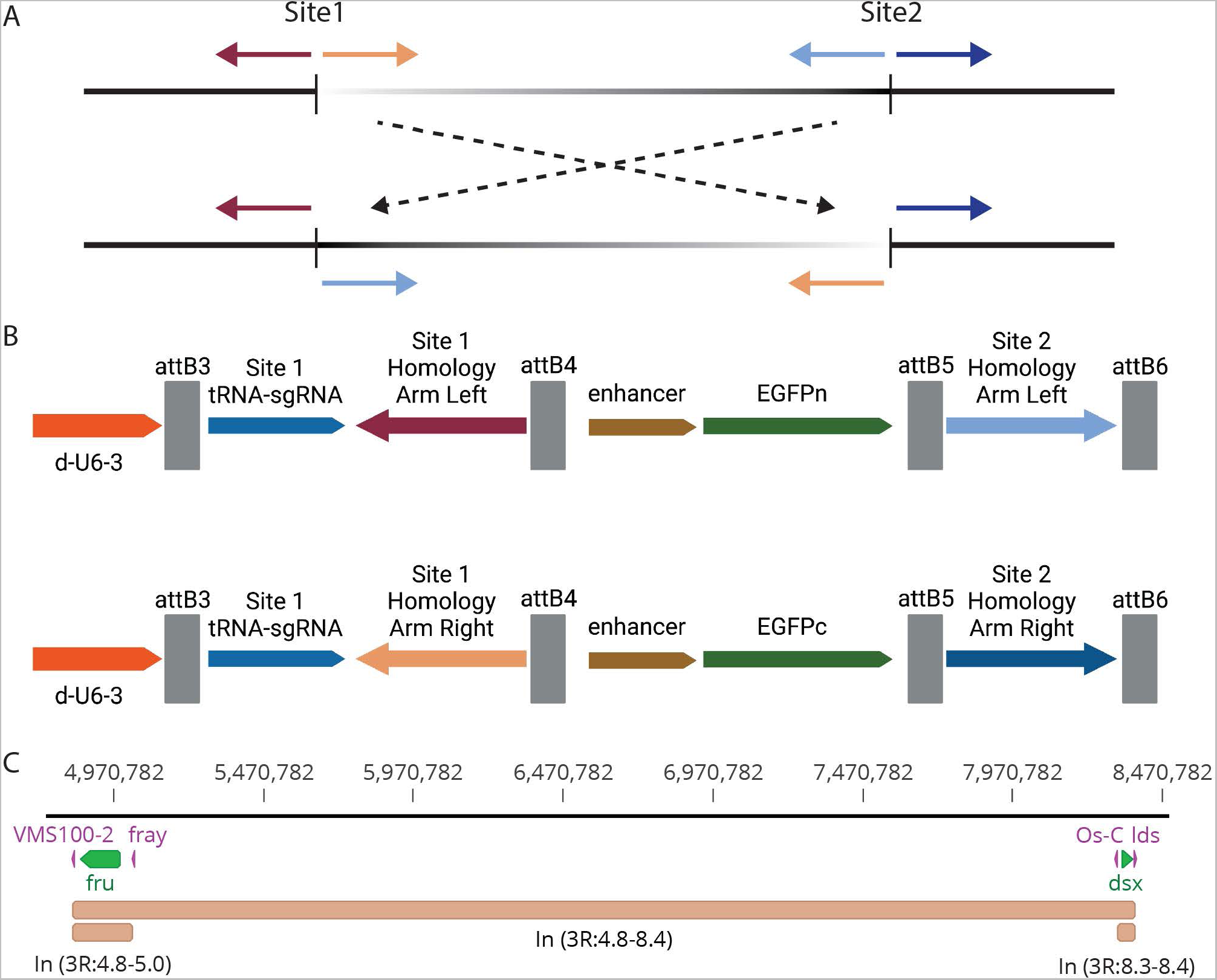
Design of pJAT plasmids for targeted inversions. (A) Schematic illustrating the position and orientation of DNA sequences flanking two cut sites before (top) and after (bottom) a chromosomal inversion. (B) Illustration of two pJAT plasmids, each containing one half of a split-GFP reporter and a gRNA targeting one of the two intended cut sites. Each plasmid includes synthesized arms oriented in the direction of the intended inversion so that homology directed repair at both sites simultaneously will tend to drive an inversion. (C) Illustration of three inversions targeted on the right arm of chromosome 3 in *D. yakuba*. The positions of the *fruitless* (*fru*) and *Doublesex* (*dsx*) genes are shown in green. The locations of target gRNA sites in the genes are shown in magenta. The three targeted inversions are shown in beige.

We targeted three inversions of approximately 58 kb to 3 Mbp in *Drosophila yakuba* to provide potential balancer chromosomes for reagents that we had previously built at the *fruitless* and *doublesex* genes in this species (Figure 5c). We targeted breaks within genes that are expected to be homozygous deleterious or lethal. We detected multiple events for each targeted inversion, at efficiencies of 8-23% (Supplementary Table 4).

We PCR amplified fragments corresponding to the expected inversion event breakpoints (Supplementary Figure 6) and confirmed that the sequences of both breakpoints were oriented as expected (Supplementary Figure 7). We crossed the three *D. yakuba* inversions to null alleles of either *dsx* or *fru*, which they were designed to balance, and found that they all balanced the targeted alleles for multiple generations. Thus, single inversions can be quickly introduced to balance specific alleles in *Drosophila* species. This technology will substantially simplify the maintenance of transgenic reagents and mutants in non-*melanogaster Drosophila* species. It is possible that multiple inversions can be introduced serially onto a single chromosome arm to generate balancer chromosomes of even greater utility^10^, and we provide three sets of split-GFP reagents that may enable these experiments (Figure 2).

## Discussion

We have developed a series of new plasmids that provide a simple cloning platform and enable high efficiency HDR in multiple *Drosophila* species. The plasmid design is flexible and allows introduction of any payload of interest. The efficiency of pJAT plasmids is so high that it is almost certainly preferable to random integration via transposable elements for most purposes and may even be more efficient than the most widely used method of transgenesis in *Drosophila* at the moment, site-specific recombination into attP sites^17^. We have reported previously that some strains and some species appear to be resistant to specific transposable elements and to *attP-attB* integration^18^. In fact, we previously tried multiple times to introduce a *nos-Cas9* transgene using transposable elements into the specific *D. simulans* strain used in this study without success, but we identified multiple independent events with a single injection of a pJAT plasmid carrying *nos-Cas9*. Thus, pJAT plasmids are probably a preferable method of transgenesis to random integration using transposable elements, especially for studies of non-*melanogaster Drosophila* species.

We designed the pJAT plasmids for ease of cloning using synthesized homology arms that incorporate tRNA-sgRNA-tRNA arrays. Because these arms are synthesized, any modifications to the native sequence can be introduced during synthesis and then integrated into the genome. We illustrated the power of this approach by including mobilization sequences of the *piggyBac* transposon into the synthesized arms. This allows efficient scarless removal of the transposon and reporter gene, leaving behind substitutions introduced into the genome (Figure 4). An additional illustration of the utility of this approach is that additional CRISPR target sites can be introduced flanking the homology arms, allowing the homology arms and payload to be linearized *in vivo* (Supplementary Figure 3). This does not appear to increase HDR efficiency using pJAT plasmids in *Drosophila* (Supplementary Figure 3), which is consistent with reports of HDR induced by zinc-finger nucleases in *Drosophila* that showed that circular donor DNA was more efficient than linearized DNA^2^. However, linearization may increase efficiency of HDR or of homology-mediated end joining in other systems^19, 20^.

We initially attempted to build the double-Gateway system using multiple *ccdB* containing cassettes, as has been demonstrated by others^8^. However, we discovered during subsequent cloning that in every case one of the two *ccdB* genes had been inactivated. This may result from the dosage sensitivity of *E. coli* to *ccdB* copy number^4^. We therefore developed a novel Gateway-like second cloning cassette that exploits the well-established strong negative selective capability of the *SacB* gene^21^. When *E. coli* are grown on high levels of sucrose, expression of the *SacB* gene causes lethality. Thus, this stabilized double-Gateway system dramatically improved cloning efficiency of two synthesized fragments containing complementary *attL* sites over a design that utilizes two cassettes both using the *ccdB* gene.

While we designed the Janelia Atalanta series to simplify cloning, we found that these plasmids also increased HDR efficiency compared with other methods that we have tested and that have been reported. Often, HDR in many genes fails^3, 6^ and, when it succeeds and has been reported efficiency is approximately 1-10% on average. Although difficult to quantify, publication bias may mean these reported efficiencies are over-estimates of true efficiencies, since we are aware of multiple examples of failed CRISPR-HDR attempts in *Drosophila* species performed by other research groups. We so far have been able to target every locus attempted and on average targeting genes or genomic regions that have non-lethal mutant phenotypes results in approximately 25% of fertile injected animals yielding correct insertion events. We performed a series of experiments to try to determine what might contribute to this increased efficiency and it appears that multiple factors contribute to higher efficiency HDR. Some of the most important factors appear to be (1) production of sgRNA from a tRNA-sgRNA-tRNA array, (2) injection into embryos with a germline source of Cas9 protein, (3) homology arms of at least 250 bp long, (4) circular plasmid DNA, and possibly (5) increasing the incubation temperature to 25°C for a time after injection.

Despite the overall high efficiency of HDR with pJAT plasmids, we identified a strikingly lower efficiency on average with plasmids targeting genes with lethal null phenotypes. A subset of these “lethal” genes did show high efficiencies, however. In limited experiments, we found no evidence that lower HDR efficiencies resulted from the specific gRNA or homology arms used, but we cannot rule out the possibility that these variables influenced efficiency. One possibility which we have not tested is that the pJAT plasmids are so efficient that when the reagents work at one targeted allele in an individual, the second allele is also hit at high frequency. This would cause injected animals to be mosaic for homozygous null mutations, which may lead to lethality of sterility of many flies carrying HDR events.

The most powerful demonstration of the high efficiency of pJAT plasmids is that they could be used to introduce two simultaneous breaks in a single chromosome and facilitate inversion of large chromosomal regions through HDR (Figure 5, Supplementary Figures 5 & 6). Three independent experiments, using two separate injection companies, yielded inversions (Supplementary Table 4). These inversions provide practical benefits, since they can be used to balance deleterious alleles and will simplify stock maintenance. It may also be possible to use pJAT plasmids to generate multiply inverted balancer chromosomes with wide applicability^16^. It has not escaped our attention that this approach may also allow reversion of naturally occurring chromosomal inversions^22^, which often become hot-spots of evolutionary divergence^23^. Such experiments may allow, for the first time, detailed recombination-based identification of loci “locked” within non-recombining inversions that have contributed to phenotypic divergence.

The pJAT plasmids were designed to provide a generic platform for introducing many variant elements easily and it is possible that pJAT plasmids may provide an improved CRISPR-HDR platform for other species. One potentially species-specific component of this first set of pJAT plasmids are the U6 promoters. We used the *D. melanogaster* U6-3 promoter for all of the experiments described in this paper and we observed similar HDR efficiencies in *D. melanogaster*, *D. simulans*, and *D. yakuba* (Figure 3), suggesting that this U6-3 promoter works well for these species. This U6 promoter could be easily swapped with a different species-specific or inducible promoter. We have therefore generated pJAT77, where cloning sites can allow any promoter to be placed on either or both sides of the Gateway cloning cassettes. Some pJAT plasmids presented in this paper (Figure 2_ contain other species-specific components that may not work efficiently in other species. For example, some of these plasmids include germ-line specific promoters and 3’UTRs that were derived from the *D. melanogaster nanos* and *vasa* genes. It is possible that these will not work efficiently in other species. But the pJAT plasmids with multiple cloning sites allow efficient cloning of new payloads with other species-specific components. It will be interesting to determine whether pJAT plasmids increase HDR efficiency in organisms other than *Drosophila*.

## Methods

### Construction of Janelia Atalanta (pJAT) plasmids

pJAT plasmids were assembled by Golden Gate cloning^24^ of PCR amplified and synthesized fragments. All plasmids reported here are available from Addgene under plasmid numbers 204289-204312.

Gateway compatible homology arms were synthesized by Twist Bioscience with Adaptors On. Because these sequences contain almost identical attL sites at either end, synthesis is inefficient and often fails with Adaptors Off. In addition, since these fragments are cloned into the pJAT plasmids using Golden Gate cloning, there is no need to remove the Twist synthesis adaptors. Golden Gate cloning was performed as follows.

|  |  |
| --- | --- |
| pJAT plasmid (~50 ng/uL) | 1 uL |
| Homology arm left (50 ng/uL) | 0.5 uL |
| Homology arm right(50 ng/uL) | 0.5 uL |
| LR Clonase II | 0.25 uL |

Reaction was incubated at 25°C in a thermocycler for 2-12 hours. Then 0.125 uL of Proteinase K (provided with LR clonase kit) was added and sample was incubated in a thermocycler at 37°C for 20 minutes. The entire reaction volume was gently mixed with at least 25 uL of Zymo Mix&Go Competent Cells thawed on ice. The mixture was incubated for 30 minutes on ice, heat shocked for 30 seconds at 42°C, and returned to ice. 100 uL of SOC was added and the mixture was shaken at 200 RPM at 37°C for 1 hour. The entire mixture was plated on Tryptone-Yeast + Sucrose plates + Spectinomycin plates. The entire protocol, including a guide for designing and ordering homology arms, is provided at Protocols.io: dx.doi.org/10.17504/protocols.io.bp2l6bjokgqe/v1.

### Injection of Drosophila embryos

Most pJAT plasmids were injected into Drosophila embryos by Rainbow Transgenics and some were injected by Genetivision. Initially, plasmids were co-injected with Cas9 RNA generated by *in vitro* transcription from plasmid encoding Cas9^10^. After we generated fly lines carrying a *nanos*-*Cas9* transgene (*pJAT17*), plasmids were injected into flies homozygous for *pJAT17* and fertile adults emerging from injected eggs were outcrossed to a wild-type strain and offspring were screened for fluorescent reporter genes. The genomic insertion location of all integrants were determined using TagMap^11^ with the following TagMap PCR primers, where the underlined sequence represents the Tn5 MEB adaptor and the non-underlined sequence represents the sequence specific to each pJAT plasmid.

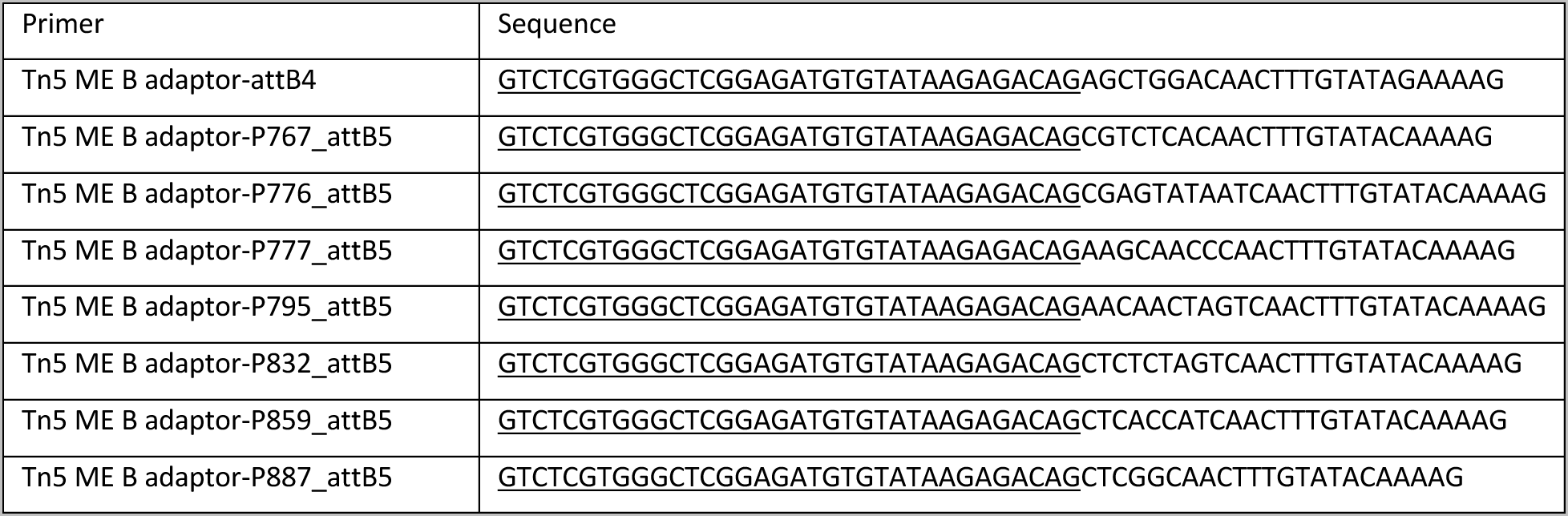

### Design and generation of scarless site specific mutations

Gateway compatible fragments were designed to include the 5’ and 3’ pBac sequences required for transposition internal to the targeting homology arms (Figure 4). Fragments were synthesized by Twist Biosciences and Gateway cloned into pJAT10 as described above.

### Design and generation of targeted inversions

Gateway compatible fragments were designed with homology arms configured to promote chromosomal inversions (Figure 5). Fragments were synthesized by Twist Biosciences and Gateway cloned into pJAT21 and pJAT25 as described above. Both plasmids were injected simultaneously into flies carrying a nos-Cas9 transgene.

## Acknowledgements

We thank Carlos Machado, Thomas Ravenscroft, and Gerry Rubin for helpful comments on the manuscript. Elements of Figures 1, 4, and 5 and Supplementary Figures 2, 3, and 4 were created with BioRender.com. We especially thank Justine Ayelet Stern for designing and creating Supplementary Figure 1 and for recommending the name Atalanta for these plasmids.

## Author Contributions

D.L.S. conceptualized, designed, and constructed most of the plasmids and performed much of the fly work. E.K. assisted with construction and quality control of plasmids and with fly work. E.L.B. constructed pJAT17 and confirmed its utility. The paper was written by D.L.S. with input from E.K. and E.L.B.

## Ethics Declarations

HHMI has filed a provisional patent, number 63/507,335, for the inventor D.L.S. covering vectors and methods for efficient cloning and homology directed repair.

## Supplementary Figure Legends

**Supplementary Figure 1.**
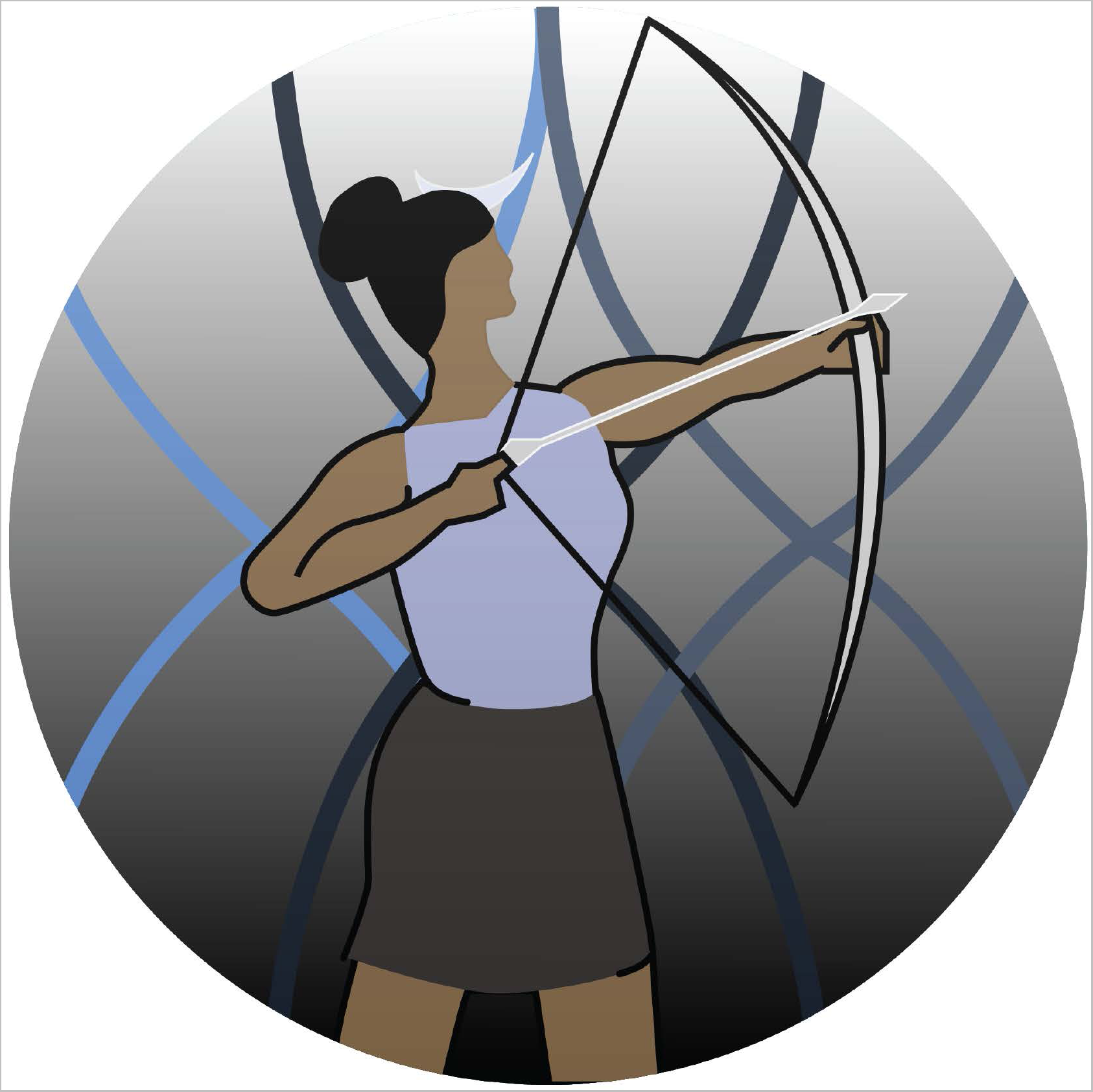
Icon for the Janelia Atalanta series of plasmids. Representation of the Greek goddess Atalanta practicing archery (artwork by Justine Ayelet Stern).

**Supplementary Figure 2.**
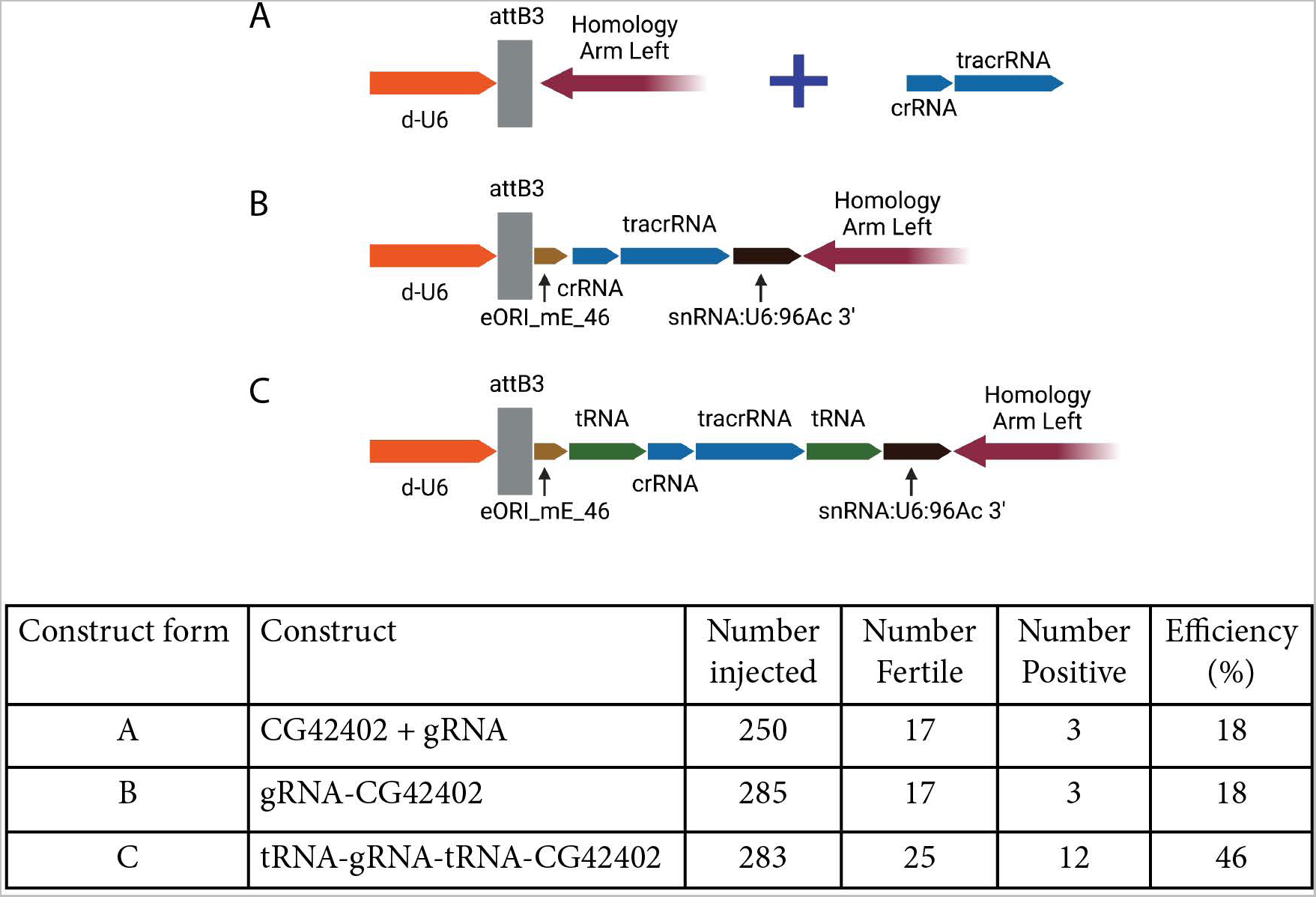
Effect of tRNA-sgRNA-tRNA array on CRISPR-Cas9 HDR efficiency. Three plasmids were created where the left homology arm contained either no tRNA and sgRNA (A), just the sgRNA (B), or the full tRNA-sgRNA-tRNA array (C). Injections of the plasmid without a sgRNA were supplemented with *in vitro* transcribed sgRNA. All plasmids targeted the *CG42402* gene. The table shows the HDR efficiency for each construct type.

**Supplementary Figure 3.**
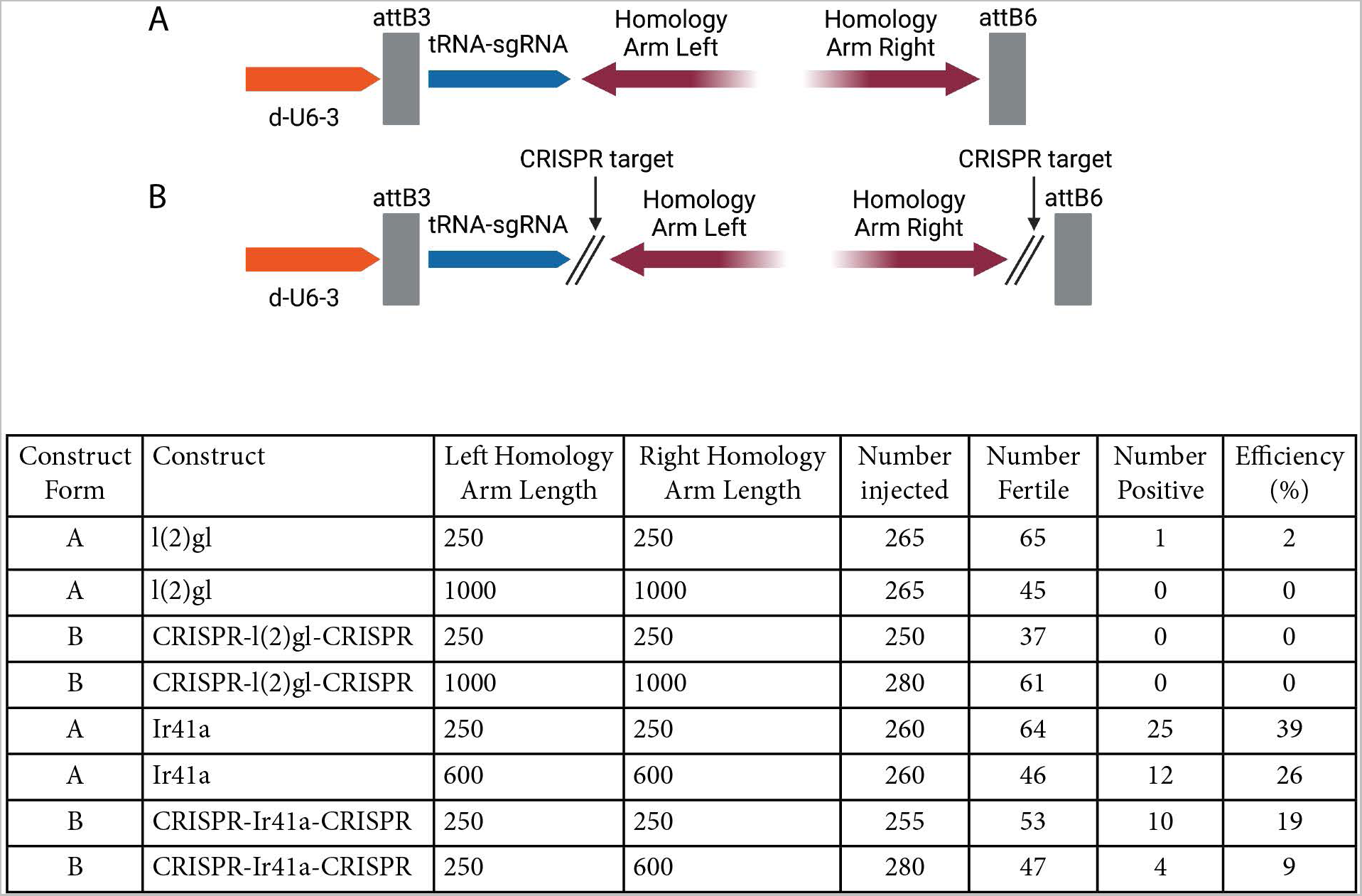
To test the hypothesis that *in vivo* linearization of the template DNA increases the efficiency of HDR, plasmids targeting two loci were constructed either without (A) or with (B) CRISPR target sites for the sgRNA encoded in the tRNA-sgRNA-tRNA array. Results are shown in the table, which illustrates that plasmids containing the flanking CRISPR sites did not increase HDR efficiency for two genes.

**Supplementary Figure 4.**
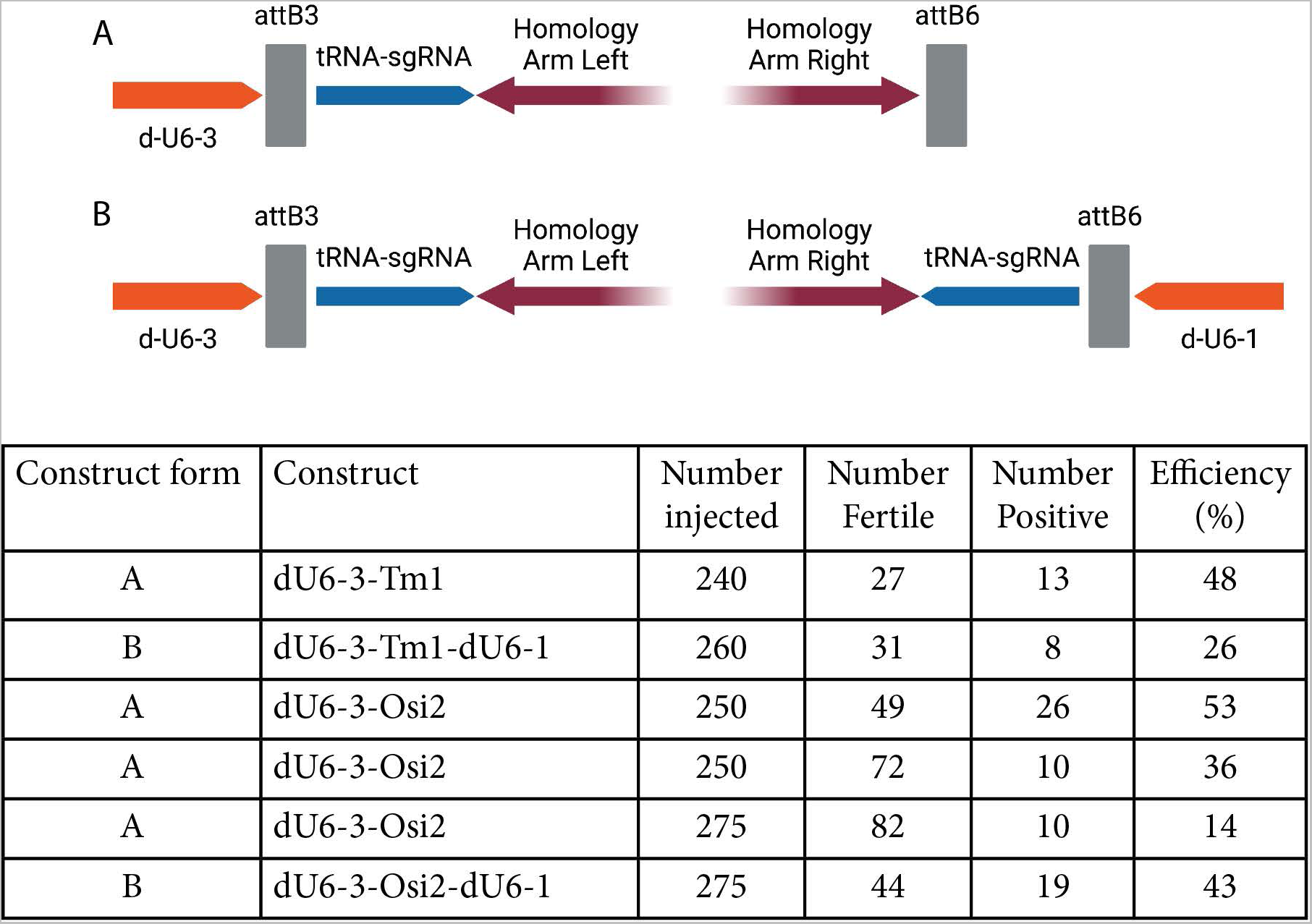
To test the hypothesis that sgRNA concentration limited HDR, plasmids including either one (A) or two (B) sources of sgRNA were constructed. Results are shown in the table, which illustrates that plasmids containing two sources of sgRNA did not increase HDR efficiency for two genes on average.

**Supplementary Figure 5.**
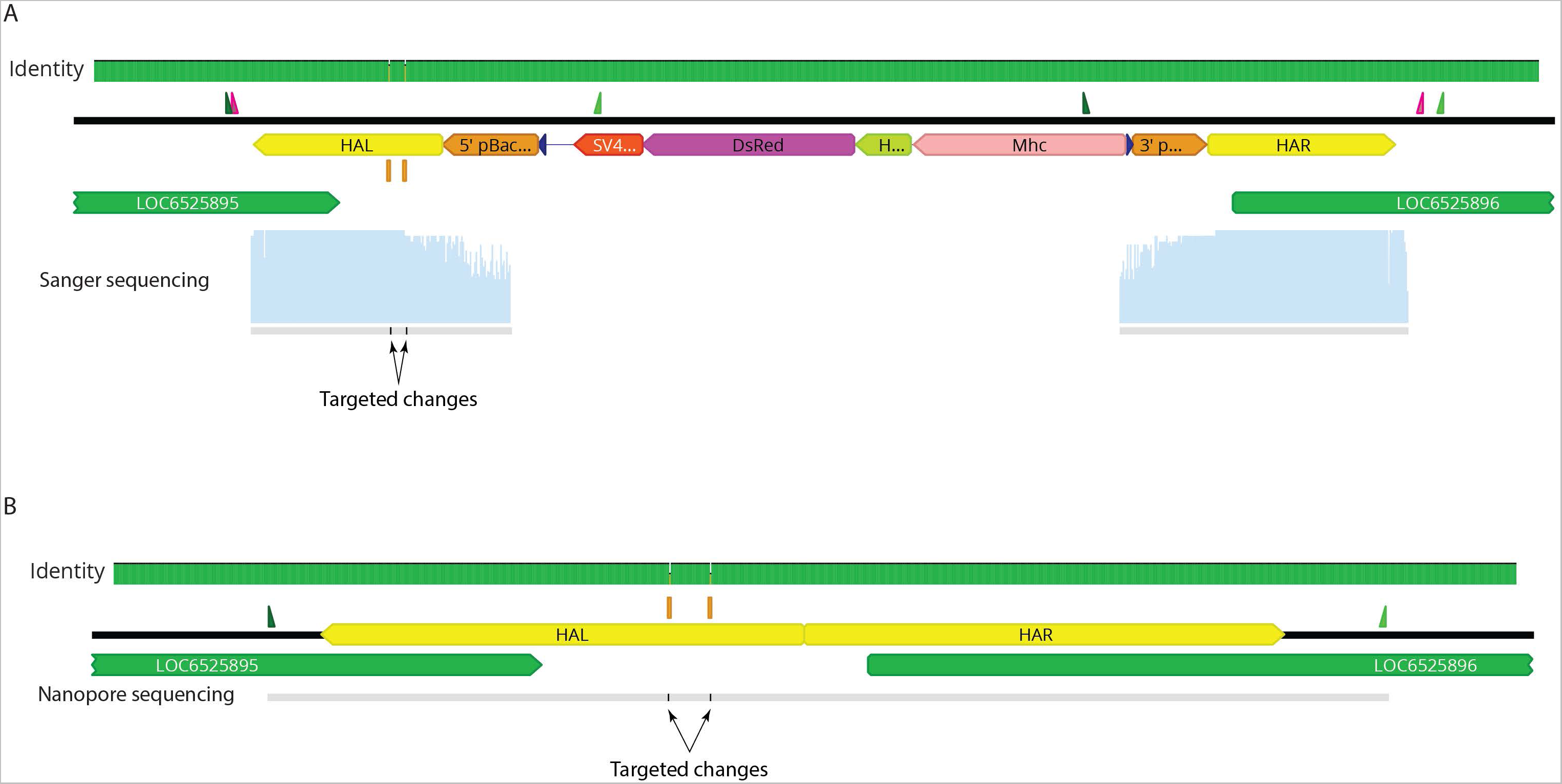
Sequence confirmation of scarless genome mutagenesis. (A) Sequencing of genomic DNA containing inserted pBac transposable element with internal MHC-DsRed reporter. Left homology arm includes intended 2 bp changes. PCR fragments were generated with primers external to homology arms (green) and internal to the pBac transposon and Sanger sequenced with internal primers (magenta). Sanger sequencing products were aligned to the original genome sequence and mismatches are shown in the Identity track and below the Sanger sequencing read. (B) A PCR product of the final scarless allele generated with primers shown (green) was Nanopore sequenced and aligned to the original genome sequence and mismatches are shown in the Identity track and within the consensus Nanopore sequencing read.

**Supplementary Figure 6.**
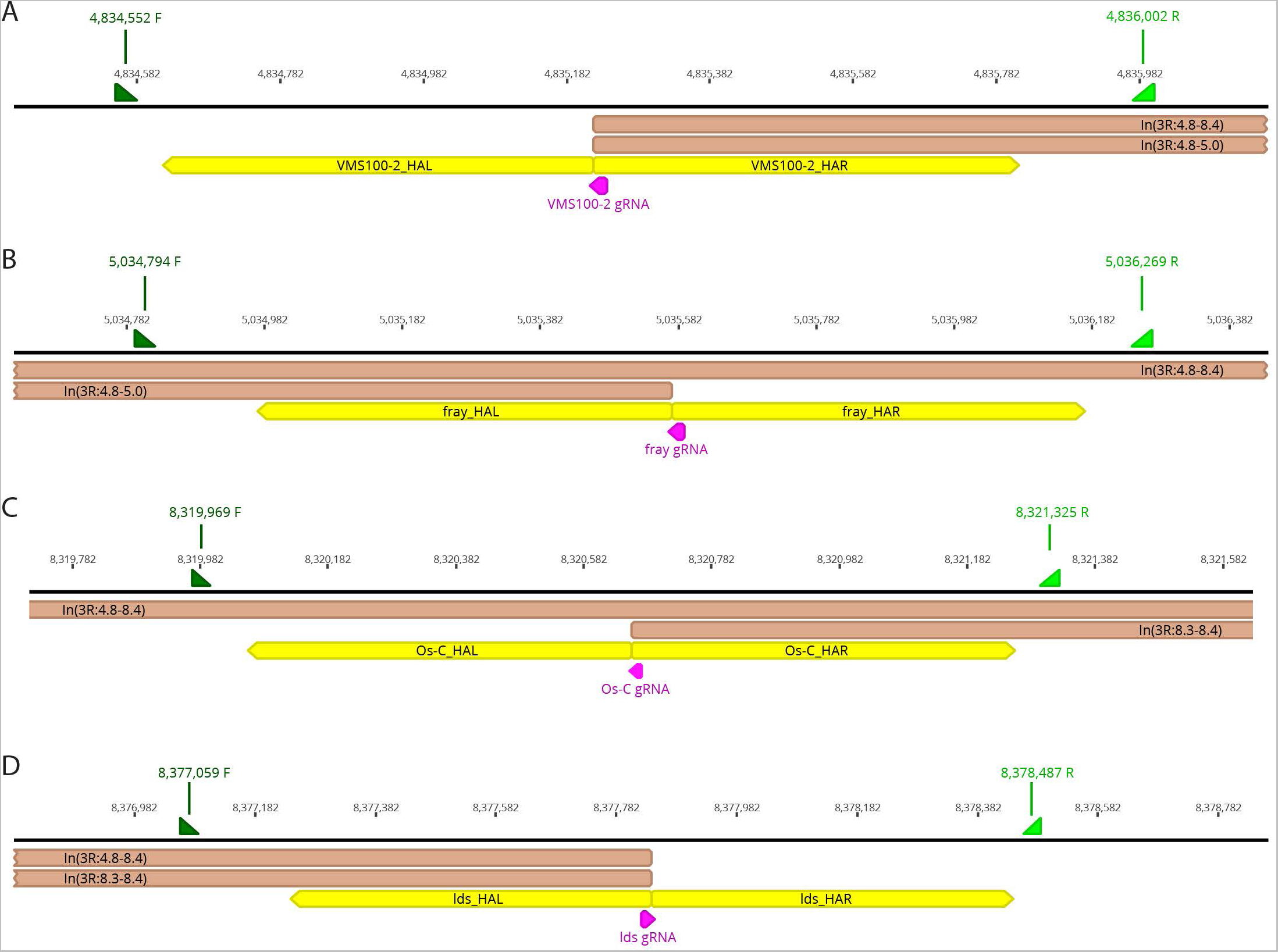
Detailed maps from the *D. yakuba* genome of the gRNA sites (magenta), homology arms (yellow), intended inversion ends (brown), and PCR primers for the genes *VMS100-2* (A), *fray* (B), *Os-C* (C), and *Ids* (D).

**Supplementary Figure 7.**
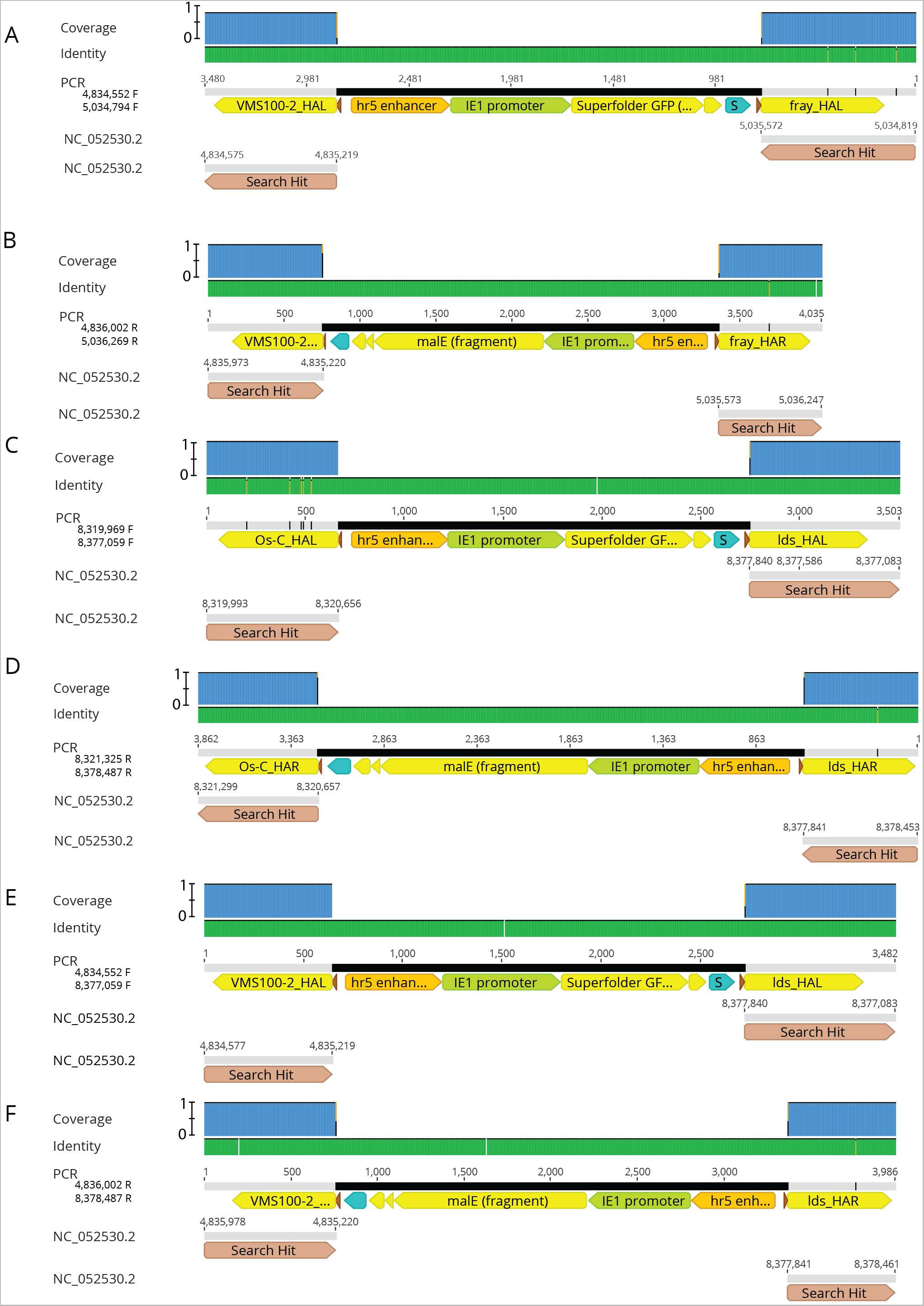
Sequences of PCR products generated using primers illustrated in Supplementary Figure 6 on strains carrying the putative inversions. Three inversions were generated, one between *VMS100-2* and *fray* (A, B), one between *Osc* and *Ids* (C, D), and one between *VMS100-2* and *Ids* (E, F).

## Notes

https://www.addgene.org/depositing/82896/

### Summary of Updates

Errors in some plasmid details illustrated in Figure 2 were identified by readers and these errors have been corrected in this version.

